# Dietary fiber intervention modulates the formation of the cardiovascular risk factor trimethylamine-N-oxide after beef consumption: MEATMARK – a randomized pilot intervention study

**DOI:** 10.1101/2024.07.08.602621

**Authors:** Melanie Haas, Beate Brandl, Klaus Neuhaus, Susanne Wudy, Karin Kleigrewe, Hans Hauner, Thomas Skurk

**Affiliations:** Technical University Munich, ZIEL Institute for Food & Health, Core Facility Human Studies, 85354 Freising, Germany; Technical University Munich, ZIEL Institute for Food & Health, Core Facility Microbiome, 85354 Freising, Germany; Bavarian Center for Biomolecular Mass Spectrometry (BayBioMS), TUM School of Life Sciences, Technical University of Munich, 85354 Freising, Germany; School of Medicine and Health, Technical University Munich, 81675 Munich, Germany; Else Kröner-Fresenius-Center of Nutritional Medicine, TUM School of Life Sciences, Technical University of Munich, 85354 Freising, Germany

**Keywords:** Food intake biomarker, trimethylamine-N-oxide, microbiome, dietary fiber, intervention study

## Abstract

The gut-microbiota-dependent metabolite trimethylamine-N-oxide (TMAO) has emerged as a potential risk factor for cardiovascular disease (CVD). Conversely, dietary fiber has been associated with a reduced risk of CVD and been proposed to exhibit beneficial effects on gut health. Considering these associations, we conducted an intervention study to investigate the influence of fiber supplementation on intestinal TMAO formation and its response after beef consumption.

Our randomized, double blind, pilot study MEATMARK included thirteen volunteers who underwent a dietary fiber and placebo intervention over two weeks. We assessed the effect of fiber supplementation on the gut microbiota and expression of the enzyme *cutC*, a key enzyme for microbial TMA formation, a precursor for TMAO. We measured the TMAO response following beef consumption after the two-week intervention. We further investigated the impact of three human single nucleotide polymorphisms (SNPs) in the expression of the hepatic enzyme *FMO3* on TMAO plasma levels, as this factor is responsible for the final oxidation step from TMA to TMAO.

Our findings indicate that dietary fiber supplementation attenuated TMAO formation after beef intake, particularly in participants with habitual lower daily meat consumption. Furthermore, we observed a significant downregulation of *cutC* expression in response to the fiber intervention, suggesting a potential mechanism to reduce plasma TMAO. Considering these findings, a high fiber, low meat diet presents a promising dietary strategy for reducing this CVD risk factor.

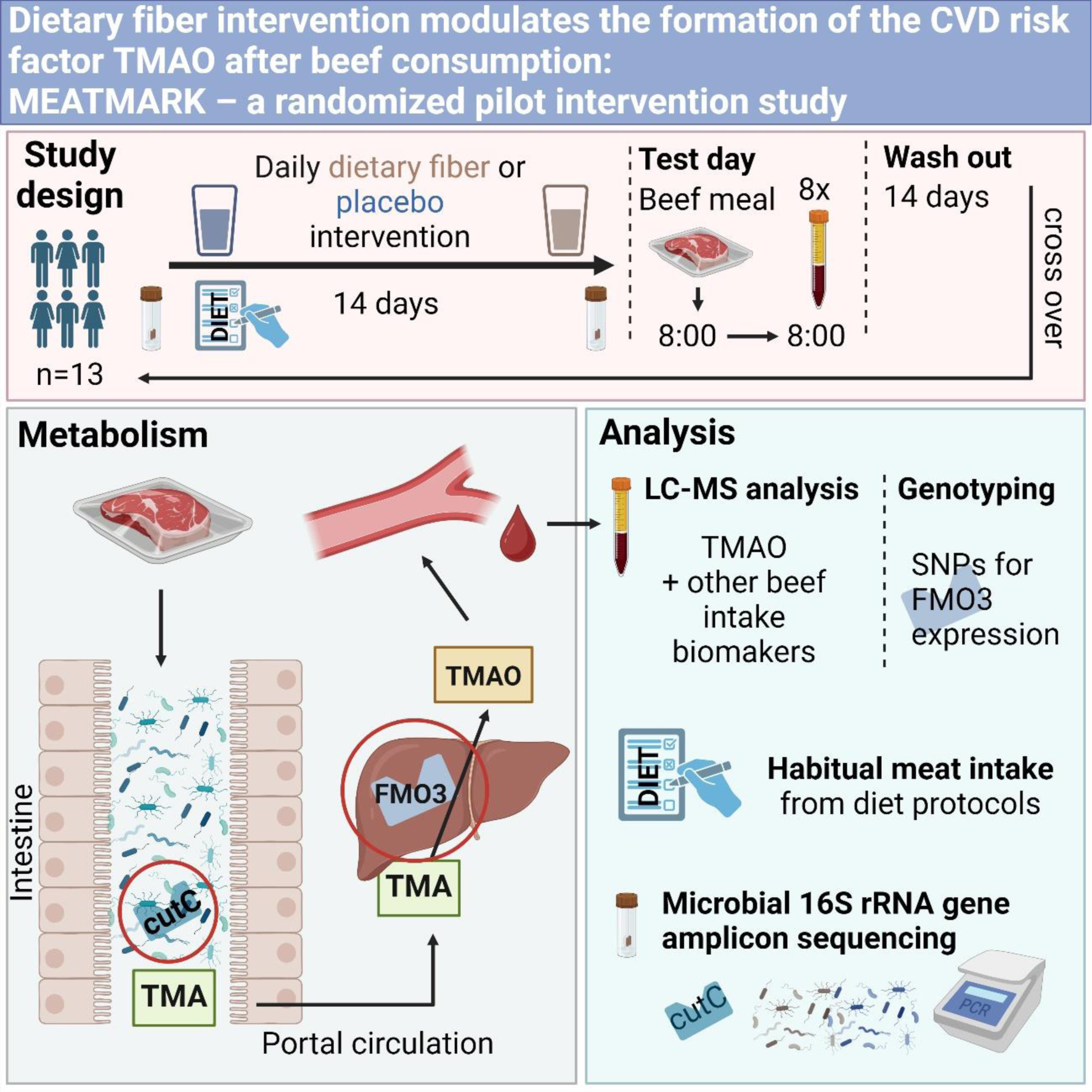

## 1. Introduction

TMAO is considered a gut microbiome-dependent biomarker and serves as a surrogate measure for meat intake and may be implicated in CVD risk ^1–4^. Recently, TMAO has garnered increasing attention in cardiovascular health research ^5,6^. Meta-analyses of epidemiological studies have shown that TMAO could be a significant risk factor for cardiovascular events ^3,4,7,8^.

Its formation is intricately influenced by diverse factors, including diet ^5,9^, age ^10^, individual microbiome composition ^5^, and the expression of specific microbial and hepatic enzymes ^11,12^. TMAO can directly originate from dietary sources like fish or seafood, but the largest proportion in the human circulation is generated stepwise via a meta-organismal pathway from dietary precursors like choline, L-carnitine and betaine ^1,13,2^. These dietary precursors can be found in various foods, like red meat (products) and eggs. Therefore, TMAO is primarily considered a food biomarker for meat and fish ^5^ After ingestion of the above mentioned dietary precursors, they are metabolized by the gut microbiome to trimethylamine (TMA) ^14,5,15^. Two main enzymatic steps are necessary for microbial TMA production: the choline TMA-lyase enzyme (*cutC*) utilizing choline, and the two-component Rieske-type oxygenase/reductase enzyme system (*cntA*) metabolizing carnitine ^16^. Subsequently, TMA is absorbed by the intestinal epithelium, transported to the liver, and is finally oxidized by flavin-containing monooxygenases (*FMOs*), mainly *FMO3* ^17^, to TMAO.

Various endogenous and exogenous factors, such as gut microbiome composition, dietary patterns, and genetic predispositions, are known to affect TMAO production and, therefore, need thorough investigation ^14^. Notably, differences in microbiome composition and basal TMAO levels already exist between individuals under a vegan and omnivore diet ^5^. In this respect, novel therapeutic strategies are being developed to modulate the microbiome or metabolic pathways in a hypothetically beneficial way ^18^. One possible approach might involve the modulation of the microbiome using dietary fibers, known for their prebiotic effects ^19,20^ and several other positive physiological effects, especially their potential role in reducing the risk of CVD ^21–23^.

Another, yet underestimated, aspect regarding TMAO and CVD could be the endogenous hepatic formation of TMAO in dependence of the host genotype. Among hepatic *FMOs*, *FMO3* plays a crucial role in producing TMAO from TMA, which is linked to trimethylaminuria, a rare genetic disorder known as “fish odor syndrome”. In the case of trimethylaminuria, mutations in the *FMO3* gene result in altered conversion of TMA to TMAO. In the case of trimethylaminuria, the genetic mutations results in a decreased expression of *FMO3* and cause accumulation of TMA in plasma resulting in body odor reminiscent of spoiled fish ^17,12^. Interestingly, trimethylaminuria does not seem to be associated with CVD or thrombosis ^12^. However, it is supposed that a deficiency in *FMO3* results in lower levels of TMAO, which is believed to reduce the risk of thrombosis and atherosclerosis.

Objectives of this investigation were to examine the interaction between a highly standardized, dietary fiber intervention, the composition of the gut microbiota, and their enzymatic and metabolic activities concerning the production of TMAO after beef consumption. As there is clear evidence that eating pattern significantly differentiates the TMAO response, we also separated the MEATMARK cohort into regular and occasional meat eaters. Additionally, to stratify endogenous TMA transformation we screened the participants for three SNPs relevant for the expression of *FMO3* (*rs2266780, rs909530* and *rs909531*).

## 2. Methods

Procedures for the intervention study were followed in accordance with the ethical standards of the Helsinki Declaration and were reviewed and approved by the ethics committee of the School of Medicine at the Technical University of Munich, Germany (331/20S). All study participants had provided written informed consent. The study was registered in the German Clinical Trials Register (DRKS): DRKS00021291.

### 2.1. Study design

The MEAT intake bioMARKer (MEATMARK) study is a six-week randomized, cross-over, double-blind pilot study involving thirteen healthy volunteers (6 females, 7 males). Each participant underwent two intervention phases, each lasting two weeks, during which they consumed either the dietary fiber supplement or a placebo made of maltodextrin (see **2.3 Study product** for details on the used products). Following each intervention phase, participants arrived fasted at the study center in the morning of the test day and consumed a test meal composed of 200 g of beef, 125 g of rice, 30 g of margarine, and 1.5 g of salt to assess any differences in TMAO metabolism after fiber supplementation compared to placebo. A two-week washout period separated the two intervention phases, as illustrated in **Figure 1**.

**Figure 1.**
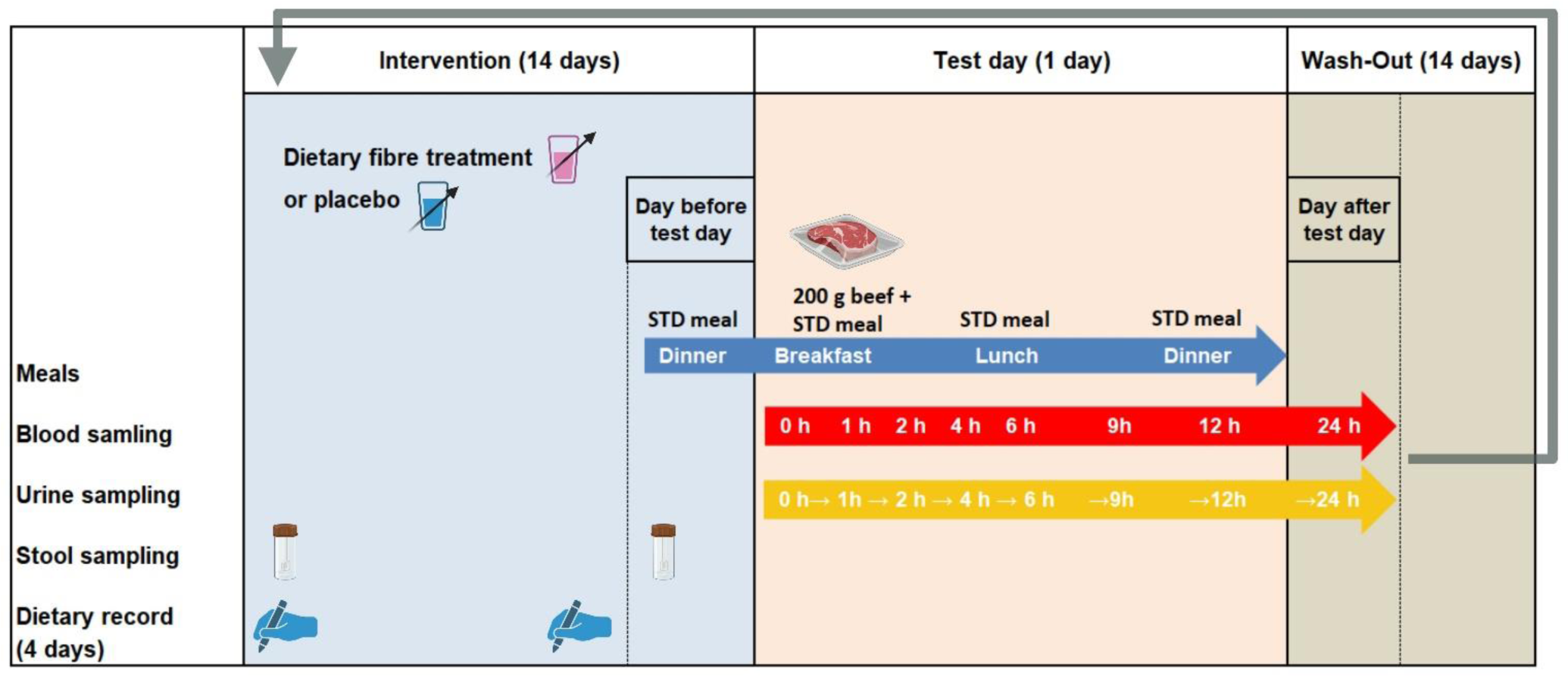
Study design of the MEATMARK study. One study cycle is shown; each participant underwent two cycles distinguished by the intervention (either dietary fiber supplement or placebo). The entire cross-over study period lasted six weeks, consisting of two two-week interventions, followed by a study day and separated by a two-week wash-out period. The STD meal was also provided the evening before test day and for lunch and dinner on test day (blue arrow). Blood (red arrow), urine (yellow arrow) and stool was sampled at defined time points and dietary records were taken on four consecutive days before and at the end of the intervention phase.

During the intervention phases, the fiber supplements, provided as powder, were suspended in water immediately prior to consumption. To get used to the high-fiber supplement, participants continuously increased their intake from 9 to 27 g of fiber or placebo over time. On the evening before a test day, participants received a standardized meal (STD meal) comprising 125 g of rice, 30 g of margarine, and 1.5 g of salt. For breakfast on intervention days, participants consumed 200 g *sous-vide* cooked beef alongside the STD meal. The STD was also provided for lunch and dinner during the course of the intervention day.

On the test day, blood and urine samples were collected at defined intervals over a 24-hour period, while stool samples were collected before and after each two-week intervention phase. Participants were instructed to document their food intake for four consecutive days before the intervention phase and for the last four days during the intervention phase and to categorize the stool according to the Bristol stool scale. Additionally, participants refrained from consuming foods containing TMAO or its precursors (including meat, fish, eggs, and foods containing L-carnitine) for two days prior to the intervention day.

### 2.2. Study participants

For this trial, 24 healthy volunteers (18–40 years) were recruited via flyers at the Technical University of Munich, campus Freising. Participant enrollment started on August 10^th^, 2020, and continued until December 1^st^, 2020. The participants’ eligibility was assessed with a detailed screening questionnaire. Exclusion criteria were age <18 years and >40 years, smoking, diseases affecting nutrient absorption, digestive, metabolic, or excretory function, chronic diseases from the medical history (e.g. hepatitis B and C, diabetes mellitus), antibiotic use in the previous six months, regular medication intake (except oral contraceptives), pregnant and breastfeeding women, known allergies or food intolerances to any of the test food ingredients, diverticulitis and/or constipation in the medical history. We enrolled 16 volunteers from which 3 were excluded for analysis because of premature termination of the study due to personal reasons or antibiotic use. Finally, 13 participants were included and **Figure 2** depicts the participant flow chart of the MEATMARK study.

**Figure 2.**
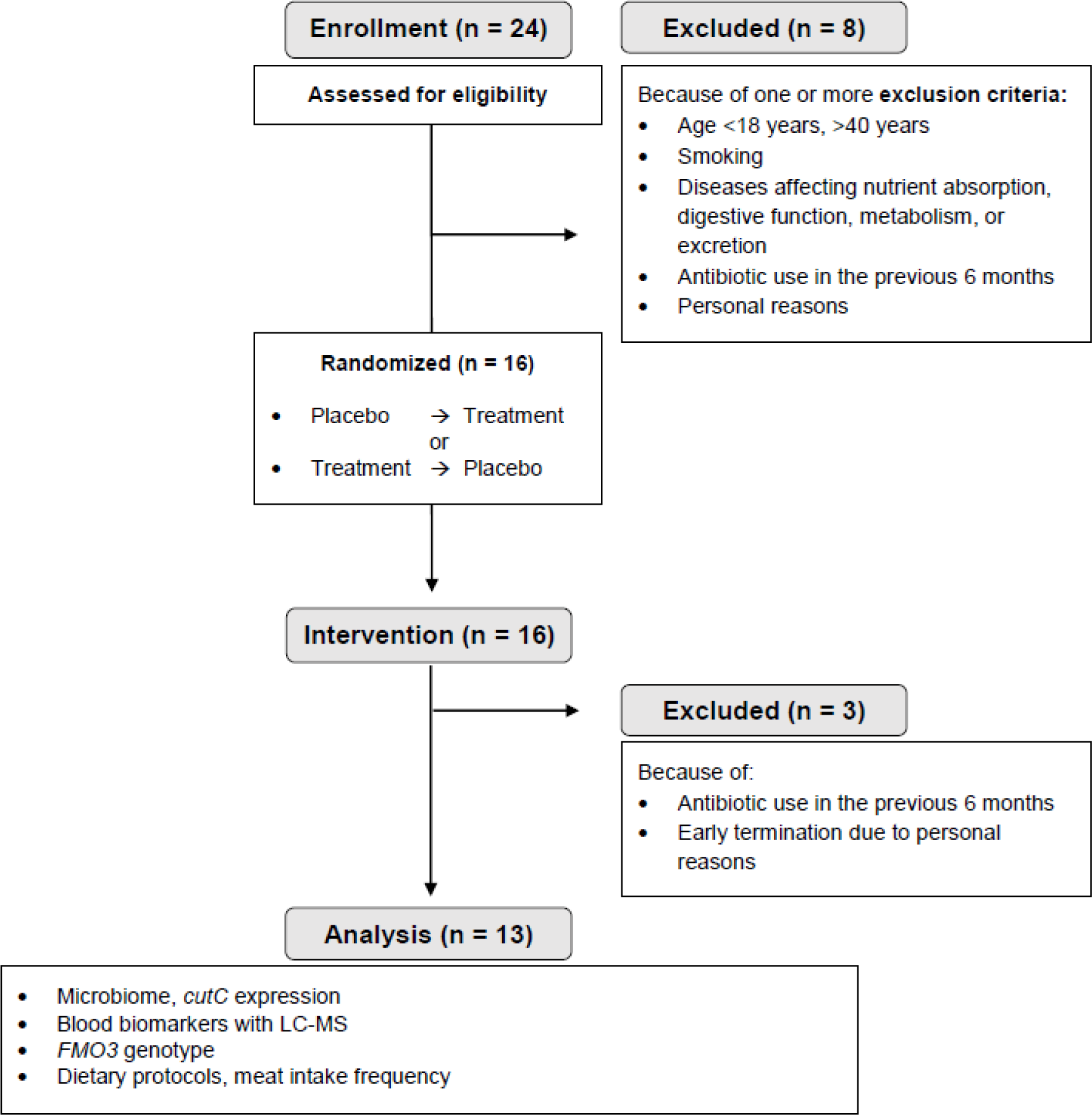
Consort flow diagram of study participants.

### 2.3. Study products

The dietary fiber supplement, as well as the placebo product, were provided by J. Rettenmaier & Söhne GmbH + Co KG, (Rosenberg, Germany). Both products were supplied in powdered form and should be suspended in water for consumption.

The dietary fiber product comprised 48% total dietary fiber, consisting of 87% wheat fiber, 7% psyllium, and 6% guar gum. The remaining components of the product included 42% isomaltulose, citric acid, sucralose, and some flavors and food colorants. Conversely, the placebo product contained a total dietary fiber content of 10%, supplemented by 25% maltodextrin, 7.5% vegetable potato flakes, 50% isomaltulose, and citric acid, sucralose, and the same flavors, and food colorants. The detailed composition is provided in the supplementary material (see **Table S1: Composition of study products** in the supplementary material).

Each product was packed in sachets of 19.05 g, corresponding to 9 g of total dietary fiber. Participants were instructed to increase their intake of dietary fiber progressively: from 9 g (days 1-2), to 18 g (days 3-6), and finally to 27 g (days 7-14). The recommended method of consumption was to dissolve the powder in 200 mL of water and to consume it throughout the day alongside a meal.

### 2.4. Blood, buffy coat and fecal sampling

#### Plasma and buffy coat sampling

Blood samples were collected in the fasting state and after 1, 2, 4, 6, 9, 12 and 24 h after the beef meal as breakfast on the test day. Plasma (EDTA K2 monovettes, Sarstedt, Nümbrecht, Germany) was collected and centrifuged at 1,800 × *g* for 10 min at 4°C. Serum was collected, allowed to clot for 30 min, and then centrifuged (2,500 × *g* for 10 min at 4°C). Buffy coat was sampled from EDTA tubes (EDTA K3 monovettes, Sarstedt) after it was centrifuged at 1,800 × *g* for 10 min at room temperature. Plasma, buffy coat and serum were aliquoted and stored at −80°C for later measurements.

#### Stool sampling

Before and at the end of every intervention phase (**Figure 1**), the participants collected stool samples in two separate tubes according to a standardized procedure. One tube contained 8 mL DNA stabilization buffer (Stratec Molecular GmbH, Berlin, Germany). Participants were instructed to bring the samples to the study center within 6 hours, where they were subsequently stored at −80°C until analysis.

### 2.5. Targeted amino acid and acylcarnitine measurement

Targeted amino acid and acylcarnitine measurement were performed using a QTRAP 5500 triple quadrupole mass spectrometer (Sciex, Darmstadt, Germany) coupled to an ExionLC AD (Sciex, Darmstadt, Germany) ultrahigh performance liquid chromatography system. A multiple reaction monitoring (MRM) method was performed based on Wudy et al. ^24^, with extension by the following analytes (see **Table S2: MRM transitions for quantification of amino acids and acylcarnitines** in the supplementary material). Briefly, for chromatographic separation of amino acids and acylcarnitines, a 2.1 × 100 mm, 100 Å, 1.7 μm, UPLC BEH amide column (Waters, Eschborn, Germany) was used with 5 mM ammonium acetate in water (eluent A) and 5 mM ammonium acetate in acetonitrile/water (95/5, v/v) (eluent B) as elution solvents both adjusted to pH 3 using acetic acid. An injection volume of 1 µL and a flow rate of 0.4 mL/min was used. The gradient elution started at 100% B and was held for 1.5 min.

Afterward, the concentration was decreased to 92% B at 3.5 min and further reduced to 90% B at 7 min. At 10 min, 78% B was used and decreased to 65% B, further decreased to 2% B at 12 min, and held for 1 min. At 15.5 min, the column was equilibrated at starting conditions. The column oven was set to 40 C and the autosampler to 15 C. Ions of amino acids and acylcarnitines were analyzed in the positive electrospray ionization mode. The electrospray voltage was set to 5,500 V, curtain gas to 35 psi, ion source gas 1 to 55 psi, ion source gas 2 to 65 psi, and the temperature to 400 C. The MRM parameters (**Table S2**) were optimized using commercially available standards. Data acquisition and instrumental control were performed with Analyst v1.7 software (Sciex, Darmstadt, Germany). The data was analyzed with MultiQuant v3.0.3 software (Sciex, Darmstadt, Germany).

### 2.6. High-Throughput 16S rRNA gene sequencing and *cutC* gene expression from microbiome DNA

#### Microbial 16S rRNA gene amplicon sequencing

The 16S rRNA gene sequencing was conducted as described in Reitmeier et al. ^25^. Briefly, DNA was isolated after bead-beating using guanidinium thiocyanate as denaturing agent and N-lauroylsarcosine for separation. Finally, polyvinylpyrrolidone was used to remove phenolic inhibitors from the DNA ^26^. After RNase A digest, 12 ng of the isolated and cleaned DNA was used for library preparation in a 2-step PCR. The first PCR uses primer for the V3-V4 region ^27^; 341F, CCTACGGGNGGCWGCAG; 785r-ovh, GACTACHVGGGTATCTAATCC) with overhangs for the second PCR. The subsequent PCR adds barcodes and adapter for Illumina sequencing. PCR products were cleaned and sequenced in equimolar batches on a MiSeq using PE300 cartridges.

Raw reads were processed using the UPARSE approach implemented in IMNGS using SOTUs as output ^28^. Spurious taxa were filtered at <0.25% abundance per sample ^29^. Denoised amplicons were assigned to taxa using Silva v138. Reads were normalized to the lowest read number of a given sample as described. Alpha and beta-diversity, taxonomic binning, as well as correlations were assessed using the pipeline Rhea, a collection of R-scripts for 16S rRNA data analysis ^30^.

#### *CutC* expression

Gene expression from the 1V-2V region of *cutC* was performed by quantitative PCR (qPCR) in a LightCycler 480 (Roche Diagnostics, Mannheim, Germany) using the Maxima SYBR Green/ROX qPCR Master Mix (ThermoFisher Scientific, Waltham, Massachusetts, USA) according to the manufacturer’s instructions. PCR settings and procedure were as follows: an initial 95°C step for 10 min was followed by 40 cycles of denaturation at 95°C for 30 s; annealing at 52°C for 45 s; and an extension step at 72 C for 45 s. Melting curves were subsequently performed using the following program: 95°C for 10 s, followed by 60°C for 1 s, and a final continuous reading step of two acquisitions per second between 60 and 95°C.

For microbial *cutC* gene expression analysis, a fold-change (FC) analysis after the intervention compared to the respective baseline was performed.The primer for *cutC* was previously described by Rath et al. ^15^ and are as follows: cutC-F, TTYGCIGGITAYCARCCNTT; cutC-R,TGNGGYTCIACRCAICCCAT.

### 2.7. DNA isolation and *FMO3* genotyping

Double-stranded DNA was isolated from buffy coat using the DNeasy Blood & Tissue Kit with DNeasy Mini spin-columns (QIAGEN, Hilden, Germany) following the manufacturer’s protocol. For genotyping, three different SNPs of the *FMO3* gene (*rs2266780, rs909530*, and *rs909531*) have been examined for mutations by using a melting curve analysis with LightSNiP assays from TIB MOLBIOL (TIB MOLBIOL, Berlin, Germany) in a LightCycler 480 (Roche Diagnostics, Mannheim, Germany). We followed the manufacturer’s protocol except for the polymerase, where we used MyTaq DNA Polymerase (Bioline, Heidelberg, Germany). The SNPs were selected after a search in the GTEx portal (https://www.gtexportal.org).

### 2.8. Dietary protocol and meat intake questionnaire

The study participants were instructed to record their food consumption before the intervention phase and at the end of the intervention phase on four consecutive days. The energy content and macronutrient composition of the diets were calculated using the OptiDiet Plus software (Version 5.1.2.046, GOE GmbH, Linden, Germany).

For the meat intake questionnaire, participants were asked how often meat and meat products were consumed on a regular basis. For analysis, report of eating meat less than three times a week was classified as ‘occasional’ (meat intake) and eating meat more than three times a week was classified as ‘regular’ (meat intake).

### 2.9. *enable* study

To address our research question further, we used data of the well-phenotyped cross-sectional enable study and performed FMO3 genotyping genotyping (see **2.7. DNA isolation and *FMO3* genotyping**). This analysis also involved data from targeted LC-MS analysis (plasma TMAO level) and results from usual intake and food frequency questionnaires (FFQ) ^31^. The phenotyping program of different age cohorts included, beyond others, anthropometry, body composition analysis, health and functional status, and assessment of dietary intake including food preferences and aversions. 459 healthy volunteers from different age groups including young adults (18-25 years; n = 94), middle-aged adults (40-65 years “middle-agers,” *n* = 205), and older adults (75-85 years; *n* = 160) underwent this program. Further details on the study design and the characteristics of the cohorts can be found elsewhere ^31,32^.

Additionally, we measured *cutC* expression in stool samples from a nested intervention study with middle-agers presenting an elevated waist circumference (>102 cm in males, >88 cm in females) indicating a higher cardiometabolic risk was used to (see **2.6. High-throughput 16S rRNA gene sequencing and *cutC* gene expression from microbiome DNA**). Samples and study design from this “Freising Fiber Acceptance Study” was recently described by Brandl et al. elsewhere ^32^. This subcohort was chosen for analysis according to the fiber supplementation included in the study similar to our MEATMARK cohort.

### 2.10. Data analysis and statistics

In the following, “treatment” corresponds to the fiber intervention and “placebo” corresponds to the control intervention.

Data were analyzed in R v3.6.0+ and GraphPad Prism v10.1.2 programming environment. Results are presented as mean ± SD (unless stated otherwise) and *p*-values < 0.05 were regarded as statistically significant (*). Shapiro–Wilk tests and quantil-quantil-plots were used to test for normal distribution of the data. According to sample size and distribution, either the (paired) *t*-test, the Wilcoxon matched-pairs signed rank test or the Mann-Whitney test were applied. For more than two groups, according to distribution an ordinary one-way ANOVA with Tukey’s multiple comparisons test or the Friedman test with Dunn’s multiple comparisons test was applied.

## 3. Results

### 3.1. Baseline characteristics of MEATMARK participants

Concerning inclusion and exclusion criteria, 24 volunteers were screened. Finally, 16 volunteers were enrolled. However, three participants dropped out during the study (**Figure 2**). Thus, the final analysis dataset included 13 participants (6 females, 7 males) with a mean age of 28.6 ± 6.4 years and a mean BMI of 23.1 ± 2.4 kg/m^2^. A more comprehensive description of baseline characteristics is provided in **Table S3: Baseline characteristics** in the supplementary material. The dietary protocol analysis of the placebo and treatment intervention phases are shown in **Table 1**. Notably, total energy and macronutrient composition remained consistent across the two interventions. As expected, there was a statistically significant increase in total daily fiber intake during the treatment phase, which was 51.7 ± 7.56 g/day compared to 28.7 ± 5.45 g/day.

**Table 1.**
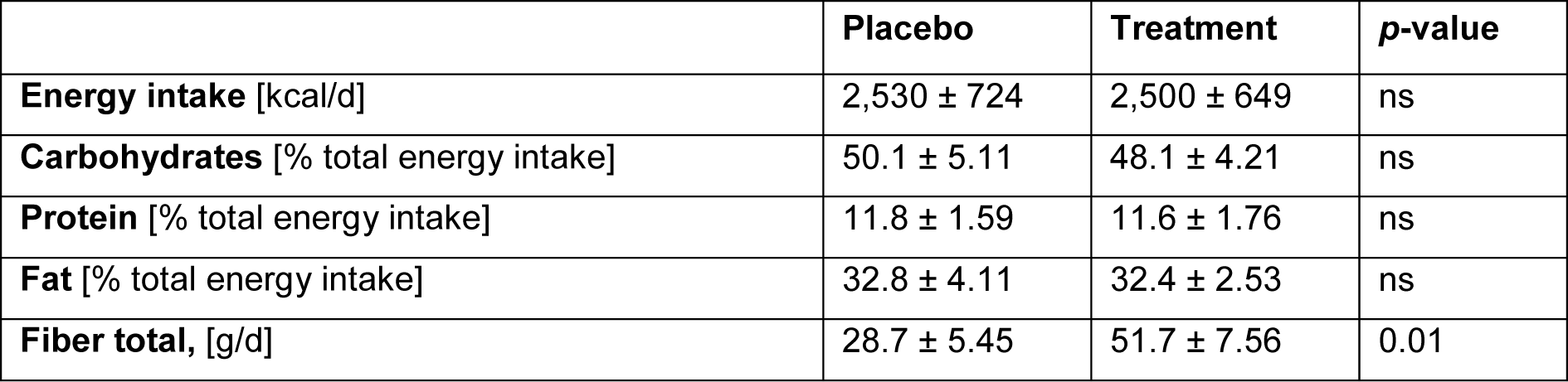
Analysis of the dietary protocol. Data is presented as mean ± SD. *P*-value < 0.05 was regarded as statistically significant; ns, not significant. According to normality distribution, either the paired *t*-test or the Wilcoxon-signed ranked test was applied to assess differences between placebo and treatment.

|  | Placebo | Treatment | *p*-value |
| --- | --- | --- | --- |
| **Energy intake** [kcal/d] | 2,530 ± 724 | 2,500 ± 649 | ns |
| **Carbohydrates** [% total energy intake] | 50.1 ± 5.11 | 48.1 ± 4.21 | ns |
| **Protein** [% total energy intake] | 11.8 ± 1.59 | 11.6 ± 1.76 | ns |
| **Fat** [% total energy intake] | 32.8 ± 4.11 | 32.4 ± 2.53 | ns |
| **Fiber total**, [g/d] | 28.7 ± 5.45 | 51.7 ± 7.56 | 0.01 |

### 3.2. TMAO as a beef intake biomarker

**Figure 3** shows plasma concentrations for creatine, 3-methylhistidine, 4-hydroxyproline and TMAO in the MEATMARK cohort after the beef intake following the 14 days fiber or placebo intervention. Creatine, 3-methylhistidine and 4-hydroxyproline are validated biomarkers of meat intake ^13,1^. These 4 analytes were indicative for beef consumption, with plasma concentrations increasing after meat consumption and returning to baseline values after 24 hours, except TMAO, which appears to remain in the blood for more than 24 hours. Pure descriptive analysis of the total group revealed no significant differences between the dietary fiber treatment and the placebo when comparing the maximum values and the AUC (see corresponding **Table S4: AUC and maximum values of beef intake biomarkers** in the supplementary material). Fold changes of TMAO baseline to individual maximum levels did also not differ significantly between interventions (*p*-value = 0.26; data in **Figure S1** in the supplementary material).

**Figure 3.**
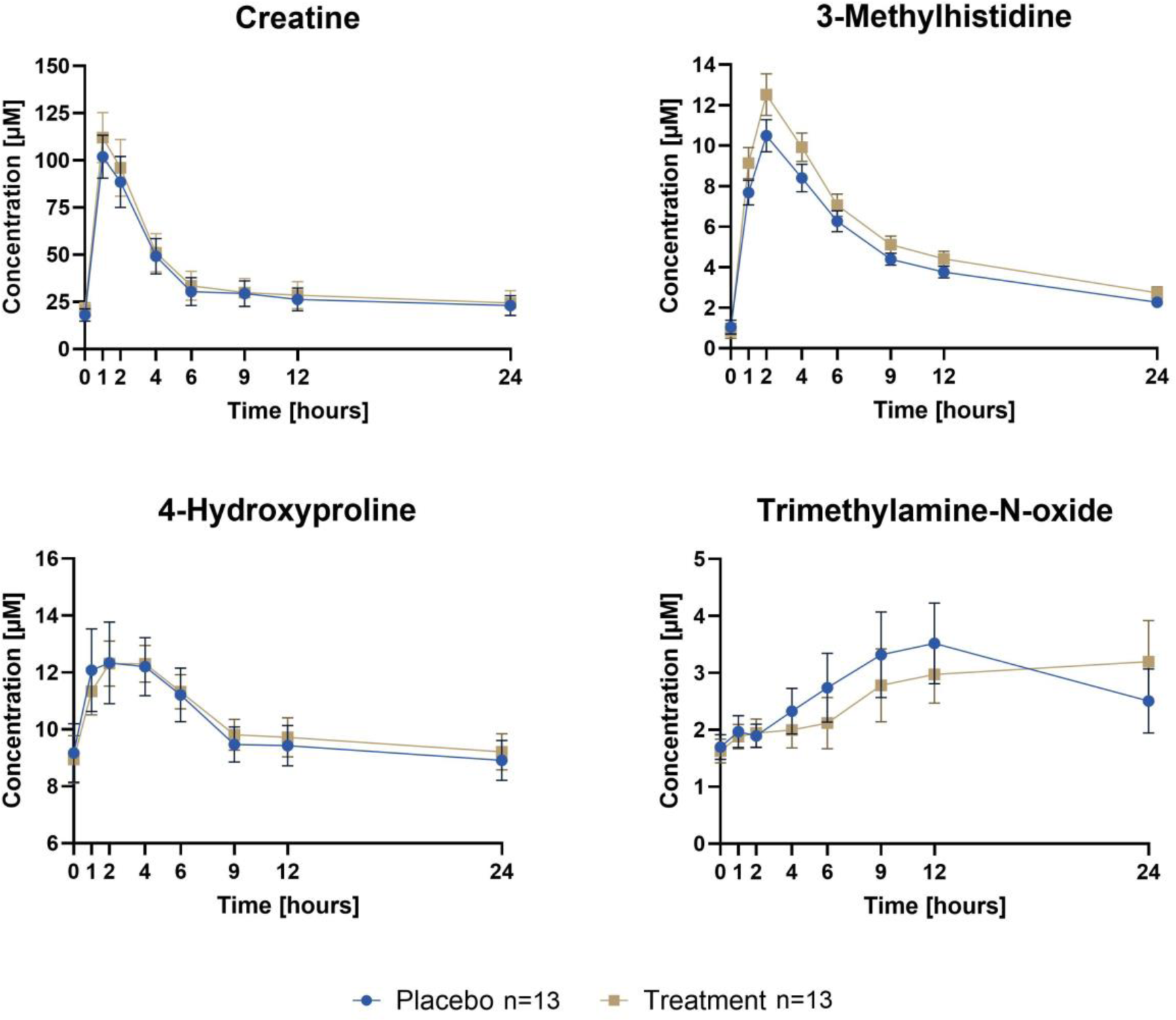
Plasma concentrations of meat intake biomarkers after beef consumption on the test day after 14 days of either fiber treatment or placebo intervention. Data is shown as mean ± SEM.

### 3.3. Occasional versus regular meat intake showed a different plasma TMAO response after two weeks of high-fiber intervention

Given the known variations in TMAO metabolism between individuals adhering to vegetarian or omnivorous diets ^5^, we first aimed to explore these differences within the *enable* cohort in more detail. As shown in **Figure 4A**, this study group revealed significant difference in basal plasma TMAO levels between vegetarians/vegans and omnivores (*p*-value = 0.028).

**Figure 4.**
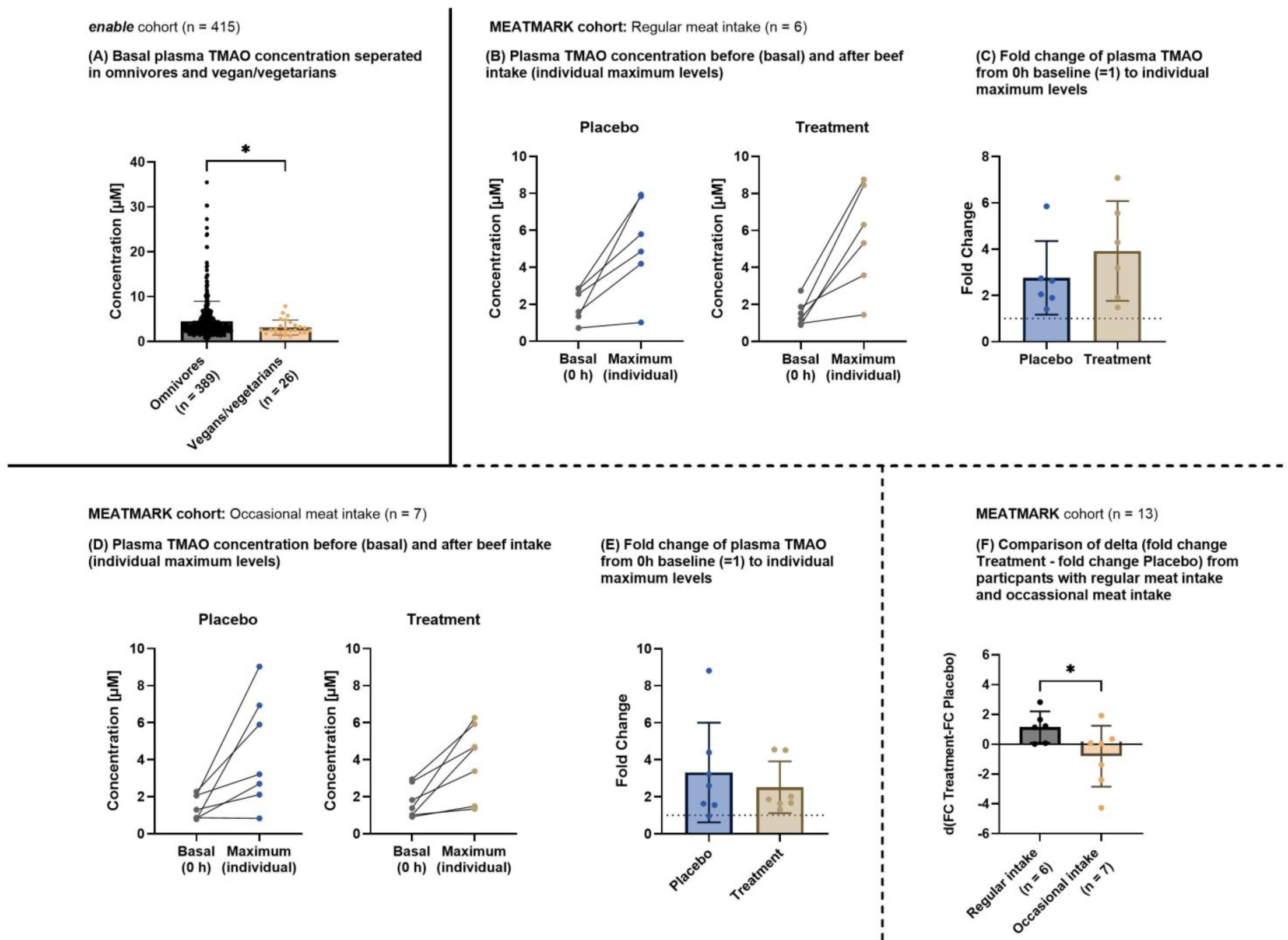
**A** Basal plasma TMAO concentrations in the *enable* cohort separated in omnivores and vegan/vegetarians. Data is presented as individual values (dots) and as mean (bar plot). According to normality distribution, the Mann-Whitney test was applied to assess differences between omnivores and vegans/vegetarians. **B** Plasma TMAO concentration before (basal) and after beef intake in the MEATMARK cohort from participants with regular meat intake. The respective individual maximum level within 24 hours is given. **C** The fold change of plasma TMAO from basal to individual maximum levels from participants with regular meat intake in the MEATMARK cohort. According to normality distribution, the Wilcoxon matched-pairs signed rank test was applied to assess differences between placebo and treatment. **D** Plasma TMAO concentrations before (basal) and after beef intake in the MEATMARK cohort from participants with occasional meat intake. The respective individual maximum level within 24 hours is shown. **E** The Fold change of plasma TMAO from basal to individual maximum levels from participants with occasional meat intake in the MEATMARK cohort. According to normality distribution, the Wilcoxon matched-pairs signed rank test was applied to assess differences between placebo and treatment. **F** Delta between FC placebo and treatment from participants with regular and occasional meat intake. According to normality distribution, the *t*-test was applied to assess differences between regular and occasional meat intake. For all panels, a *p*-value < 0.05 was regarded as statistically significant.

Although we were not able to confirm this effect in the MEATMARK cohort due to the small sample size (*p*-value = 0.27, data not shown), we further stratified this group according to meat-eating frequency. Volunteers reporting meat consumption more than three times per week were classified as “regular” meat eaters, while those consuming meat three times or less were categorized as “occasional” meat eaters. Basal (i.e., before the beef test meal) and the individual maximum plasma TMAO values (i.e., after beef consumption) for both groups were compared after treatment and placebo interventions (**Figure 4B/D**). Interestingly, while maximum TMAO values appeared comparable between regular and occasional meat eaters after the placebo phase, a notable discrepancy emerged after the fiber intervention. Specifically, the increase in maximum TMAO values after beef consumption was less pronounced in the occasional meat-eater group compared to the regular meat-eater group. This effect was further elucidated by examining the FC of TMAO values (from basal to maximum value) within each intervention group (**Figure 4C/E**). Notably, in the occasional meat eater group, the FC of TMAO decreased after the treatment intervention, in contrast to the observed increase in the regular meat eater group.

Furthermore, a comparative analysis of the delta effect of the FC between placebo and treatment revealed a significant difference between the regular and occasional meat eater groups (**Figure 4F**). The occasional meat eater group exhibited a significantly greater reduction in TMAO levels after the fiber intervention compared to the regular meat-eating participants (*p*-value = 0.029).

These findings underscore the differential response in TMAO levels to dietary fibers among individuals with varying meat consumption patterns, suggesting a potential approach for targeted dietary strategies to modulate TMAO metabolism.

### 3.4. Dietary fiber intervention alters expression of the TMA-producing microbial enzyme *cutC*

The analysis of the microbiome unveiled significant inter-individual differences in beta diversity (PERMANOVA *p*-value_gUnifrac_ < 0.001), indicating that microbial compositions among individuals differed noticeably (**Figure 5A**). With regard to our primary research question of whether a fiber intervention affects the microbiome, we analyzed and compared the samples from the four sampling times (before and after placebo and treatment intervention). The average individual richness at baseline placebo was 55 ± 17 operational taxonomic units at species level (SOTUs), 60 ± 12 SOTUs at baseline treatment, 55 ± 14 SOTUs after placebo and 55 ± 17 SOTUs after treatment. The Shannon effective number of species was 30 ± 9 at baseline placebo, 34 ± 9 at baseline treatment, 30 ± 10 after placebo, and 31 ± 10 after treatment. Both, richness and Simpson effective numbers did not differ significantly between sampling times. Beta-diversity between sampling times also showed no significant difference, respectively no significant clusters were discernible based on the sampling times (**Figure 5B**). The relative abundances of dominant gut microbiome phyla also remained remarkably consistent across all sampling times. **Figure 5C** illustrates the relative abundance of gut microbiota at family level at baseline and after placebo and treatment intervention, with *Lachnospiraceae*, *Ruminococcaceae*, *Bacteroidaceae*, *Prevotellaceae*, *Oscillospiraceae*, *Bifidobacteriaceae* and *Rikenelleceae* emerging as the predominant families. Between the two baseline samples, *Lachnospiraceae* was significant higher in the placebo baseline (*p*-value*_Lachnospiraceae_* = 0.046). *Ruminococcaceae* was significant higher after treatment intervention compared to placebo intervention (*p*-value*_Ruminococcaceae_* = 0.017) and *Bacteroidaceae* significantly increased compared to respective baseline after placebo intervention (*p*-value*_Bacteroidaceae_* = 0.037).

**Figure 5.**
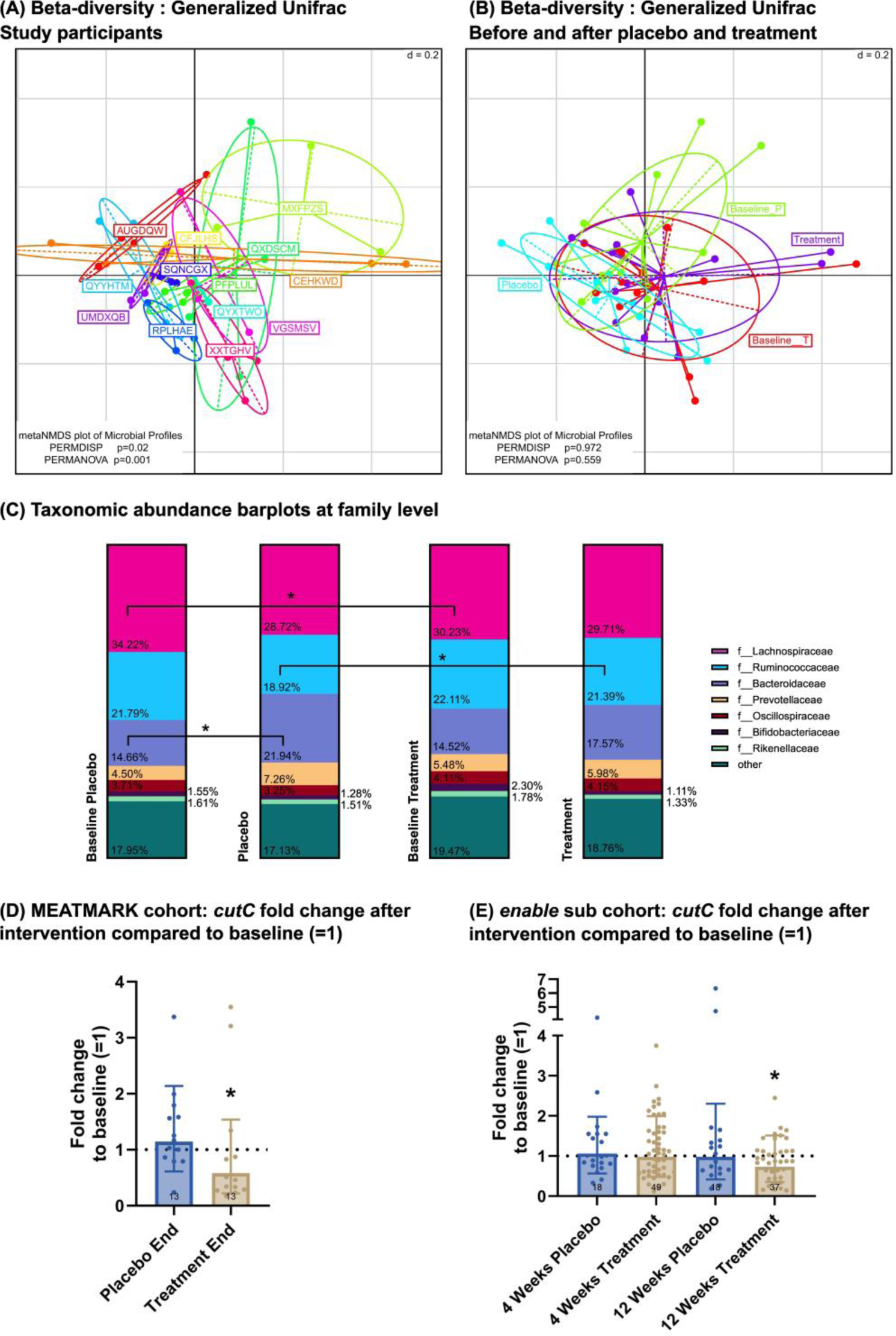
Fecal microbiota analysis by 16S rRNA gene amplicon analysis. **A** Beta diversity of individuals (every color represents one individual volunteer) and **B** at baseline and after two weeks of intervention with either placebo or treatment; PERMDISP and PERMANOVA were applied as statistical test. **C** Taxonomic bar plots on family level with relative abundance at baseline and after two weeks of placebo and treatment intervention. Data is presented as mean. According to normality distribution, either a one-way ANOVA with Tukey’s multiple comparisons test or the Friedman test with Dunn’s multiple comparisons test was applied. **D/E** Fold change of microbial *cutC* after placebo and treatment intervention compared to baseline fold change (=1). According to normality distribution, either the one tailed paired *t*-test or one-tailed Wilcoxon matched-pairs signed rank test was applied to assess differences between fold change after intervention and baseline. The *enable* subgroup refers to the participants from the “Freising Fiber Acceptance Study”. For all panels, a *p*-value < 0.05 was regarded as statistically significant.

Further analyses at the enzymatic level showed a change in the expression of the enzyme *cutC* expression after the treatment intervention. Specifically, there was a significant decrease in *cutC* expression after two-week treatment compared to the respective baseline in MEATMARK (*p*-value = 0.033; **Figure 5D**). In the *enable* sub-cohort “Freising Fiber Acceptance Study” ^32^, a fiber intervention was conducted for four and twelve weeks, including a placebo. A significant decrease in *cutC* expression after twelve weeks of dietary fiber intervention (*p*-value = 0.016) was found (**Figure 5E)**, but no change was observed after placebo. Based in these findings, we performed a correlation analysis and found two OTUs, which may be related to an increase in cutC expression. SOTU582 belongs to family *Lachnospiraceae* and SOUT411 belongs to family *Oscillospiraceae*, both indicate a higher cutC expression (SOTU582: Pearsońs R: 0.868, p-value < 0.0001; SOTU411: Pearsońs R: 0.743, p-value < 0.0001). **Figure S2** in the supplementary material depicts the correlation plots.

### 3.5. Investigation of three FMO3 SNPs on plasma TMAO levels

Exploring the influence of various SNPs (*rs909530, rs909531*, *rs2266780*), no conclusive evidence for an association of the three specific genotypes with TMAO plasma levels was obtained. This analysis encompassed both the MEATMARK and *enable* ^31^ cohorts, with detailed results provided in **Table 2**.

**Table 2.** Basal and maximum TMAO plasma levels according to the FMO3 SNP variation are shown. According to normality distribution, the Wilcoxon-signed ranked test was applied to assess differences in TMAO levels between meat intake frequency and sex. *P*-value < 0.05 was regarded as statistically significant. MEATMARK: in all three SNPs the wild type and heterozygous were statistically compared with a Wilcoxon-sign rank test. *enable*: For the statistical analysis, a Kruskal-Wallis multiple comparison test was used. Since none of the variations showed a significant difference, the *p-*values represents multiple comparison analysis. ns, not significant.

| **FMO3 SNP variation** | ***rs909530*** |  |  | ***p*-value** |
| --- | --- | --- | --- | --- |
|  | ***Wild type (CC)*** | ***Heterozygous (CT)*** | ***Mutation (TT)*** |  |
| **MEATMARK, n** | 6 | 6 | 1 |  |
| Mean: basal TMAO levels [μM] | 1.77 ± 0.57 | 1.47 ± 0.72 | 2.19 | ns |
| Placebo: maximum TMAO levels [μM] | 4.97 ± 2.61 | 4.11 ± 2.89 | 7.84 | ns |
| Treatment: maximum TMAO levels [μM] | 4.86 ± 1.20 | 4.00 ± 3.11 | 8.45 | ns |
| ***enable*, n** | 155 | 87 | 112 |  |
| Basal TMAO levels [μM] | 3.25 ± 2.39 | 4.02 ± 4.46 | 3.92 ± 2.88 | ns |
|  | ***rs909531*** |  |  |  |
|  | ***Wild type (TT)*** | ***Heterozygous (TC)*** | ***Mutation (CC)*** |  |
| **MEATMARK, n** | 8 | 3 | 0 |  |
| Mean: basal TMAO levels [μM] | 1.54 ± 0.62 | 1.42 ± 0.27 | - | ns |
| Placebo: maximum TMAO levels [μM] | 4.33 ± 3.20 | 5.68 ± 2.37 | - | ns |
| Treatment: maximum TMAO levels [μM] | 4.60 ± 2.45 | 4.14 ± 2.45 | - | ns |
| ***enable*, n** | 168 | 79 | 7 |  |
| Basal TMAO levels [μM] | 3.24 ± 2.31 | 4.13 ± 4.67 | 4.29 ± 3.31 | ns |
|  | ***rs2266780*** |  |  |  |

|  | *Wildtype (AA)* | *Heterozygous (AG)* | *Mutation (GG)* |  |
| --- | --- | --- | --- | --- |
| **MEATMARK**, n | 10 | 3 | 0 |  |
| Mean: basal TMAO levels [μM] | 1.74 ± 0.70 | 1.42 ± 0.27 | - | ns |
| Placebo: maximum TMAO levels [μM] | 4.53 ± 2.87 | 5.68 ± 2.37 | - | ns |
| Treatment: maximum TMAO levels [μM] | 4.91 ± 2.57 | 4.14 ± 2.45 | - | ns |
| **enable**, n | 169 | 78 | 7 |  |
| Basal TMAO levels [μM] | 3.22 ± 2.30 | 4.16 ± 4.70 | 4.50 ± 3.52 | ns |

## 4. Discussion

The precise origin and regulation of TMAO in response to nutrition is only poorly understood. With the results from the MEATAMRK study, we can provide a significant contribution to the effect of endogenous and exogenous factors on TMAO formation. First, we confirmed TMAO as a biomarker of beef consumption. The fiber intervention was performed to investigate whether it can modify TMAO plasma concentrations after beef consumption. This was not observed for the whole MEATMARK study cohort. However, when we divided the group into “occasional” and “regular” meat-eaters, we saw a lower increase of TMAO levels in the “occasional” meat-eater subgroup. In an additional analysis, we found significantly lower plasma TMAO levels between omnivores and vegetarians from the *enable* cohort in the fasting state (30). This observation was in agreement with previous reports from the literature ^5^. It is important to mention that the participants in the *enable* cohort, other than in MEATMARK, had no restrictions in their eating behavior on the days before the blood samples were taken. Consequently, TMAO precursors might also be derived from nutritional sources like fish, but also non-animal sources, including beetroot, eggs, and mushrooms ^1,2,9^.

At present, TMAO is controversially discussed as cardiovascular risk factor ^9^. In this context, the modification of plasma TMAO levels by dietary factors and/or dietary patterns may provide a plausible explanation. Recent studies have emphasized the considerable role of dietary factors for cardiovascular morbidity and mortality ^33,34^. There are many explanations how dietary factors may negatively affect cardiovascular health. A high dietary intake of saturated fat and sodium as well as a low fiber intake may be particularly relevant. It was also shown that a high meat consumption, and particularly processed meat are associated with increased cardiovascular ^35^.

The results of the MEATMARK suggest that meat consumption strongly promotes the formation of TMAO. After stratifying the participants into “regular” and “occasional” meat-eaters, we found that pretreatment with dietary fiber has a mitigating effect on postprandial TMAO formation after a meat load only in individuals with occasional meat intake compared to regular meat-eaters. This is in line with a recent report by Koeth et al. that people with lower meat consumption produce less TMAO after meat intake. They concluded that vegans and vegetarians probably have less capacity to produce TMAO ^5^. Our findings support this hypothesis, and we speculate that conditioning of the microbiome for TMAO formation is taking place under the habitual diet. Thereby, a dietary strategy that promotes reduced meat consumption and increased dietary fiber intake may mitigate TMAO production and, potentially, the associated cardiovascular risk.

The microbiome analysis also revealed promising evidence of dietary fiber’s beneficial influence on TMAO formation. Dietary fiber are attributed various health-promoting effects such as maintenance of immune homeostasis, effect on metabolic processes, glucose homeostasis, intestinal barrier integrity, and appetite regulation ^19^. All of these effects are primarily based on the fiber’s prebiotic effects. Prebiotics in general are defined as substrates that are “selectively utilized by host microorganisms conferring a health benefit” ^36^. Given that dietary fibers have a positive effect on the microbiome, we sought to determine its functional impact on TMA formation. Our findings demonstrated some microbiome changes at family level following both placebo and dietary fiber interventions. However, the impact of each observed change remains open. The ingredients of the placebo drink may have favored other bacteria in addition to the dietary fiber. In mice, it has been found that even only the food processing changed the microbiome ^37^. This indicates that the microbiome is capable of rapid adaptation ^38^. And, although Bacillota, Actinomycetota, and Pseudomonadota are the most dominant bacterial phyla which are associated with TMA formation, not every bacterium of a distinct phylum or even a genus exhibits the ability to produce it ^19,18,11^. Regarding TMAO metabolism, no bacterial family can explicitly be assigned to TMA formation, as different species within one bacterial family have the enzymatic ability for TMA formation ^15,11^. Therefore, we investigated the expression of the enzyme *cutC*, responsible for choline-derived TMA formation ^5^. Remarkably, in the MEATMARK cohort, global expression of *cutC* was significantly less after the two weeks of dietary fiber intervention, but not after the placebo intervention. Similar results were found in the *enable* cohort examined here ^32^ after twelve weeks of dietary fiber intervention. In the MEATMARK cohort, a correlation analysis also identified two SOTUs that presumably play a role in *cutC* expression. One SOTU is from the family of *Lachnospiraceae* and the other from the family of *Oscillospiraceae*. This demonstrates the potential of dietary fibers modulating microbial pathways, which affects TMAO metabolism. This underscores the beneficial role of dietary fibers. At this point, more in-depth analysis could be conducted on the two SOTUs found here, or the microbiome analysis could be deepened with regard to meat consumption.

The strength of our intervention study on the effect of fiber intake on TMAO formation is that findings are supported by the replication in another independent cohort of the *enable* cluster ^32^. This is important to note. As our study sample was small and had the character of a pilot study. Our approach of identifying a specific genotype that influences TMAO levels due to *FMO3* expression was not clearly detectable in the study cohorts. Little is known about cardiovascular diseases associated with specific *FMO3* genotypes. Our MEATMARK cohort was too small to identify genotype associations and in the enable cohort, subjects had no dietary restrictions the day before blood sampling, which can strongly influence basal TMAO levels. Therefore, analysis in larger cohorts with appropriate study designs is warranted. It is interesting to note in this context that a study by Miao et al. suggests an association between *FMO3* expression and metabolic dysfunctions, like hyperglycaemia, hyperlipidemia and atherosclerosis ^39^. Mechanistic mouse-studies also showed that the suppression of *FMO3* with antisense oligonucleotides decreases TMAO levels, thereby inhibiting both atherosclerosis and tissue sterol metabolism ^39^.

In conclusion, in the MEATMARK study, we were able to demonstrate that 27 g of dietary fiber supplementing a regular diet modulates the formation of plasma TMAO, but only in participants with a meat intake less than three portions per week. Furthermore, dietary fiber seems to have a notable impact on the microbiome regarding *cutC* expression. The results might therefore suggest that a low-meat diet combined with a high fiber intake is a promising strategy for reducing the formation of TMAO, thereby potentially contributing to cardiovascular risk reduction by a high fiber intake. However, there is a need for additional studies that further elaborate the role of TMAO for explaining the beneficial effect of high fiber intake on cardiometabolic disease risk.

## Acknowledgements

We would like to thank all the participants for their dedicated contribution in the study. We thank Margot Maier and Irmgard Sperrer for great support with monitoring data, and her technical assistance. Additionally, we are thankful to Björn Schröder for supporting us with his excellent expertise regarding microbiome. We would also like to thank Tim Englbrecht for his outstanding support in analyzing the LC-MS data.

## Author Contribution

Conceptualization, Melanie Haas and Thomas Skurk. Data curation, Melanie Haas. Funding acquisition, Thomas Skurk. Project administration, Melanie Haas. Investigation and analysis, Melanie Haas, Karin Kleigrewe and Susanne Wudy. Resources, Klaus Neuhaus, Thomas Skurk, Beate Brandl, Hans Hauner. Supervision, Thomas Skurk. Visualization and writing original draft, Melanie Haas. Review & editing, Beate Brandl, Susanne Wudy, Karin Kleigrewe, Klaus Neuhaus, Hans Hauner and Thomas Skurk.

## Highlights

- **Dietary Patterns and TMAO Levels**: We showed a significantly lower plasma TMAO levels in vegetarians compared to omnivores, suggesting that habitual dietary patterns, including lower meat consumption, influence TMAO production capacity and potential cardiovascular risk.
- **Dietary Fiber and the Microbiome:** Dietary fiber intake resulted in a significant reduction in *cutC* expression indicating that dietary fiber can beneficially impact the microbiome and reduce TMAO formation.
- **Potential of Fiber intake on Cardiovascular Health:** The findings imply that a diet low in meat and high in fiber may reduce TMAO production and an associated cardiovascular risk.

## List of Abbreviations

AUC: Area under the curve
TMAO: Trimethylamine-N-oxide
TMA: Trimethylamine
*FMO3*: Flavin-containing mono-oxygenase 3
*cutC*: Choline trimethylamine-lyase
STD: meal Standardized meal
CVD: Cardiovascular diseases
MS: Mass spectrometry
SNP: Single nucleotide polymorphism
FC: Fold change

## References

1. Subramaniam S, Fletcher C. Trimethylamine N-oxide: breathe new life. Br J Pharmacol. 2018;175:1344–1353. doi: 10.1111/bph.13959.

2. Janeiro MH, Ramírez MJ, Milagro FI, Martínez JA, Solas M. Implication of Trimethylamine N-Oxide (TMAO) in Disease: Potential Biomarker or New Therapeutic Target. Nutrients. 2018;10. doi: 10.3390/nu10101398.

3. Guasti L, Galliazzo S, Molaro M, Visconti E, Pennella B, Gaudio GV, Lupi A, Grandi AM, Squizzato A. TMAO as a biomarker of cardiovascular events: a systematic review and meta-analysis. Intern Emerg Med. 2021;16:201–207. doi: 10.1007/s11739-020-02470-5.

4. Qi J, You T, Li J, Pan T, Xiang L, Han Y, Zhu L. Circulating trimethylamine N-oxide and the risk of cardiovascular diseases: a systematic review and meta-analysis of 11 prospective cohort studies. J Cell Mol Med. 2018;22:185–194. doi: 10.1111/jcmm.13307.

5. Koeth RA, Wang Z, Levison BS, Buffa JA, Org E, Sheehy BT, Britt EB, Fu X, Wu Y, Li L, et al. Intestinal microbiota metabolism of L-carnitine, a nutrient in red meat, promotes atherosclerosis. Nat Med. 2013;19:576–585. doi: 10.1038/nm.3145.

6. Tang WHW, Wang Z, Levison BS, Koeth RA, Britt EB, Fu X, Wu Y, Hazen SL. Intestinal microbial metabolism of phosphatidylcholine and cardiovascular risk. N Engl J Med. 2013;368:1575–1584. doi: 10.1056/NEJMoa1109400.

7. Schiattarella GG, Sannino A, Toscano E, Giugliano G, Gargiulo G, Franzone A, Trimarco B, Esposito G, Perrino C. Gut microbe-generated metabolite trimethylamine-N-oxide as cardiovascular risk biomarker: a systematic review and dose-response meta-analysis. Eur Heart J. 2017;38:2948–2956. doi: 10.1093/eurheartj/ehx342.

8. Zuo K, Liu X, Wang P, Jiao J, Han C, Liu Z, Yin X, Li J, Yang X. Metagenomic data-mining reveals enrichment of trimethylamine-N-oxide synthesis in gut microbiome in atrial fibrillation patients. BMC Genomics. 2020;21:526. doi: 10.1186/s12864-020-06944-w.

9. Costabile G, Vetrani C, Bozzetto L, Giacco R, Bresciani L, Del Rio D, Vitale M, Della Pepa G, Brighenti F, Riccardi G, et al. Plasma TMAO increase after healthy diets: results from 2 randomized controlled trials with dietary fish, polyphenols, and whole-grain cereals. Am J Clin Nutr. 2021;114:1342–1350. doi: 10.1093/ajcn/nqab188.

10. Giesbertz P, Brandl B, Volkert D, Hauner H, Skurk T. Age-related metabolite profiles and their relation to clinical outcomes in young adults, middle-aged individuals, and older people. FASEB J. 2023;37:e22968. doi: 10.1096/fj.202101930R.

11. Rath S, Rud T, Pieper DH, Vital M. Potential TMA-Producing Bacteria Are Ubiquitously Found in Mammalia. Front Microbiol. 2019;10:2966. doi: 10.3389/fmicb.2019.02966.

12. Zhu W, Buffa JA, Wang Z, Warrier M, Schugar R, Shih DM, Gupta N, Gregory JC, Org E, Fu X, et al. Flavin monooxygenase 3, the host hepatic enzyme in the metaorganismal trimethylamine N-oxide-generating pathway, modulates platelet responsiveness and thrombosis risk. J Thromb Haemost. 2018;16:1857–1872. doi: 10.1111/jth.14234.

13. Giesbertz P, Brandl B, Lee Y-M, Hauner H, Daniel H, Skurk T. Specificity, Dose Dependency, and Kinetics of Markers of Chicken and Beef Intake Using Targeted Quantitative LC-MS/MS: A Human Intervention Trial. Mol Nutr Food Res. 2020;64:e1900921. doi: 10.1002/mnfr.201900921.

14. Tang WHW, Li DY, Hazen SL. Dietary metabolism, the gut microbiome, and heart failure. Nat Rev Cardiol. 2019;16:137–154. doi: 10.1038/s41569-018-0108-7.

15. Rath S, Heidrich B, Pieper DH, Vital M. Uncovering the trimethylamine-producing bacteria of the human gut microbiota. Microbiome. 2017;5:54. doi: 10.1186/s40168-017-0271-9.

16. Craciun S, Balskus EP. Microbial conversion of choline to trimethylamine requires a glycyl radical enzyme. Proc Natl Acad Sci U S A. 2012;109:21307–21312. doi: 10.1073/pnas.1215689109.

17. Dolphin CT, Janmohamed A, Smith RL, Shephard EA, Phillips IR. Missense mutation in flavin-containing mono-oxygenase 3 gene, FMO3, underlies fish-odour syndrome. Nat Genet. 1997;17:491–494. doi: 10.1038/ng1297-491.

18. Huang R, Yan L, Lei Y. The Gut Microbial-Derived Metabolite Trimethylamine N-Oxide and Atrial Fibrillation: Relationships, Mechanisms, and Therapeutic Strategies. Clin Interv Aging. 2021;16:1975–1986. doi: 10.2147/CIA.S339590.

19. Vinelli V, Biscotti P, Martini D, Del Bo’ C, Marino M, Meroño T, Nikoloudaki O, Calabrese FM, Turroni S, Taverniti V, et al. Effects of Dietary Fibers on Short-Chain Fatty Acids and Gut Microbiota Composition in Healthy Adults: A Systematic Review. Nutrients. 2022;14. doi: 10.3390/nu14132559.

20. So D, Whelan K, Rossi M, Morrison M, Holtmann G, Kelly JT, Shanahan ER, Staudacher HM, Campbell KL. Dietary fiber intervention on gut microbiota composition in healthy adults: a systematic review and meta-analysis. Am J Clin Nutr. 2018;107:965–983. doi: 10.1093/ajcn/nqy041.

21. Wanders AJ, van den Borne JJGC, Graaf C de, Hulshof T, Jonathan MC, Kristensen M, Mars M, Schols HA, Feskens EJM. Effects of dietary fibre on subjective appetite, energy intake and body weight: a systematic review of randomized controlled trials. Obes Rev. 2011;12:724–739. doi: 10.1111/j.1467-789X.2011.00895.x.

22. Weickert MO, Mohlig M, Koebnick C, Holst JJ, Namsolleck P, Ristow M, Osterhoff M, Rochlitz H, Rudovich N, Spranger J, et al. Impact of cereal fibre on glucose-regulating factors. Diabetologia. 2005;48:2343–2353. doi: 10.1007/s00125-005-1941-x.

23. Sánchez-Muniz FJ. Dietary fibre and cardiovascular health. Nutr Hosp. 2012;27:31–45. doi: 10.1590/S0212-16112012000100005.

24. Wudy SI, Mittermeier-Klessinger VK, Dunkel A, Kleigrewe K, Ensenauer R, Dawid C, Hofmann TF. High-Throughput Analysis of Underivatized Amino Acids and Acylcarnitines in Infant Serum: A Micromethod Based on Stable Isotope Dilution Targeted HILIC-ESI-MS/MS. J Agric Food Chem. 2023;71:8633–8647. doi: 10.1021/acs.jafc.3c00962.

25. Reitmeier S, Kiessling S, Clavel T, List M, Almeida EL, Ghosh TS, Neuhaus K, Grallert H, Linseisen J, Skurk T, et al. Arrhythmic Gut Microbiome Signatures Predict Risk of Type 2 Diabetes. Cell Host Microbe. 2020;28:258–272.e6. doi: 10.1016/j.chom.2020.06.004.

26. Godon JJ, Zumstein E, Dabert P, Habouzit F, Moletta R. Molecular microbial diversity of an anaerobic digestor as determined by small-subunit rDNA sequence analysis. Appl Environ Microbiol. 1997;63:2802–2813. doi: 10.1128/aem.63.7.2802-2813.1997.

27. Klindworth A, Pruesse E, Schweer T, Peplies J, Quast C, Horn M, Glöckner FO. Evaluation of general 16S ribosomal RNA gene PCR primers for classical and next-generation sequencing-based diversity studies. Nucleic Acids Res. 2013;41:e1. doi: 10.1093/nar/gks808.

28. Lagkouvardos I, Joseph D, Kapfhammer M, Giritli S, Horn M, Haller D, Clavel T. IMNGS: A comprehensive open resource of processed 16S rRNA microbial profiles for ecology and diversity studies. Sci Rep. 2016;6:33721. doi: 10.1038/srep33721.

29. Reitmeier S, Hitch TCA, Treichel N, Fikas N, Hausmann B, Ramer-Tait AE, Neuhaus K, Berry D, Haller D, Lagkouvardos I, et al. Handling of spurious sequences affects the outcome of high-throughput 16S rRNA gene amplicon profiling. ISME Commun. 2021;1:31. doi: 10.1038/s43705-021-00033-z.

30. Lagkouvardos I, Fischer S, Kumar N, Clavel T. Rhea: a transparent and modular R pipeline for microbial profiling based on 16S rRNA gene amplicons. PeerJ. 2017;5:e2836. doi: 10.7717/peerj.2836.

31. Brandl B, Skurk T, Rennekamp R, Hannink A, Kiesswetter E, Freiherr J, Ihsen S, Roosen J, Klingenspor M, Haller D, et al. A Phenotyping Platform to Characterize Healthy Individuals Across Four Stages of Life - The Enable Study. Front Nutr. 2020;7:582387. doi: 10.3389/fnut.2020.582387.

32. Brandl B, Rennekamp R, Reitmeier S, Pietrynik K, Dirndorfer S, Haller D, Hofmann T, Skurk T, Hauner H. Offering Fiber-Enriched Foods Increases Fiber Intake in Adults With or Without Cardiometabolic Risk: A Randomized Controlled Trial. Front Nutr. 2022;9:816299. doi: 10.3389/fnut.2022.816299.

33. Meier T, Gräfe K, Senn F, Sur P, Stangl GI, Dawczynski C, März W, Kleber ME, Lorkowski S. Cardiovascular mortality attributable to dietary risk factors in 51 countries in the WHO European Region from 1990 to 2016: a systematic analysis of the Global Burden of Disease Study. Eur J Epidemiol. 2019;34:37–55. doi: 10.1007/s10654-018-0473-x.

34. Mozaffarian D. Dietary and Policy Priorities for Cardiovascular Disease, Diabetes, and Obesity: A Comprehensive Review. Circulation. 2016;133:187–225. doi: 10.1161/CIRCULATIONAHA.115.018585.

35. Rohrmann S, Linseisen J. Processed meat: the real villain? Proc Nutr Soc. 2016;75:233–241. doi: 10.1017/S0029665115004255.

36. Gibson GR, Hutkins R, Sanders ME, Prescott SL, Reimer RA, Salminen SJ, Scott K, Stanton C, Swanson KS, Cani PD, et al. Expert consensus document: The International Scientific Association for Probiotics and Prebiotics (ISAPP) consensus statement on the definition and scope of prebiotics. Nat Rev Gastroenterol Hepatol. 2017;14:491–502. doi: 10.1038/nrgastro.2017.75.

37. Wenderlein J, Böswald LF, Ulrich S, Kienzle E, Neuhaus K, Lagkouvardos I, Zenner C, Straubinger RK. Processing Matters in Nutrient-Matched Laboratory Diets for Mice-Microbiome. Animals (Basel). 2021;11:862. doi: 10.3390/ani11030862.

38. David LA, Maurice CF, Carmody RN, Gootenberg DB, Button JE, Wolfe BE, Ling AV, Devlin AS, Varma Y, Fischbach MA, et al. Diet rapidly and reproducibly alters the human gut microbiome. Nature. 2014;505:559–563. doi: 10.1038/nature12820.

39. Miao J, Ling AV, Manthena PV, Gearing ME, Graham MJ, Crooke RM, Croce KJ, Esquejo RM, Clish CB, Vicent D, et al. Flavin-containing monooxygenase 3 as a potential player in diabetes-associated atherosclerosis. Nat Commun. 2015;6:6498. doi: 10.1038/ncomms7498.

